# Foundation Species Across a Latitudinal Gradient in China

**DOI:** 10.1101/2020.03.15.986182

**Authors:** Xiujuan Qiao, Jiaxin Zhang, Yaozhan Xu, Xiangcheng Mi, Min Cao, Wanhui Ye, Guangze Jin, Zhanqing Hao, Xugao Wang, Xihua Wang, Songyan Tian, Xiankun Li, Wusheng Xiang, Yankun Liu, Yingnan Shao, Kun Xu, Weiguo Sang, Fuping Zeng, Mingxi Jiang, Haibao Ren, Aaron M. Ellison

## Abstract

Foundation species play important roles in structuring forest communities and ecosystems. Foundation species are difficult to identify without long-term observations or experiments and their foundational roles rarely are identified before they are declining or threatened. We used new statistical criteria based on size-frequency distributions, species diversity, and spatial codispersion among woody plants to identify potential (“candidate”) foundation species in 12 large forest dynamics plots spanning 26 degrees of latitude in China. We used these data to identify a suite of candidate foundation species in Chinese forests; test the hypothesis that foundation woody plant species are more frequent in the temperate zone than in the tropics; and compare these results with comparable data from the Americas to suggest candidate foundation genera in Northern Hemisphere forests. We identified more candidate foundation species in temperate plots than in subtropical or tropical plots, and this relationship was independent of the latitudinal gradient in overall species richness. Two species of *Acer*, the canopy tree *Acer ukurunduense* and the shrubby treelet *Acer barbinerve* were the only two species that met both criteria in full to be considered as candidate foundation species. When we relaxed the diversity criteria, *Acer, Tilia*, and *Juglans* spp., and *Corlyus mandshurica* were frequently identified as candidate foundation species. In tropical plots, the tree *Mezzettiopsis creaghii* and the shrubs or treelets *Aporusa yunnanensis* and *Ficus hispida* had some characteristics associated with foundation species. Species diversity of co-occurring woody species was negatively associated with basal area of candidate foundation species more frequently at 5- and 10-m spatial grains (scale) than at a 20-m grain. Conversely, Bray-Curtis dissimilarity was positively associated with basal area of candidate foundation species more frequently at 5-m than at 10- or 20-m grains. Our data support the hypothesis that foundation species should be more common in temperate than in tropical or boreal forests, and suggest that in the Northern Hemisphere that *Acer* be investigated further as a foundation tree genus.

## Introduction

A foundation species is a single species (or a group of functionally similar taxa) that dominates an assemblage numerically and in overall size (e.g., mass or area occupied), determines the diversity of associated taxa through non-trophic interactions, and modulates fluxes of nutrients and energy at multiple control points in the ecosystem it defines (Ellison, 2019). Because foundation species are common and abundant, they generally receive less attention from conservation biologists, conservation professionals, and natural-resource managers who emphasize the study, management or protection of rare, threatened, or endangered species (Gaston and Fuller, 2007, 2008). However, protecting foundation species before they decline to non-functional levels can maintain habitat integrity, thereby protecting associated rare species at lower cost and less effort (Ellison and Degrassi, 2017; Degrassi et al., 2019).

Identifying foundation species is difficult because it can take many years—often decades— to collect enough data to distinguish foundation species from other species that also are common, abundant, or dominant (*sensu* Grime, 1987) but lack “foundational” characteristics (Baiser et al., 2013; Ellison, 2014, 2019). Rather than investigating one common or dominant species at a time in myriad ecosystems, Ellison and his colleagues have worked with data from individual and multiple large forest dynamics plots within the ForestGEO network^1^ to develop statistical criteria that can suggest which tree species might merit further attention as candidate foundation species in forests (Buckley et al., 2016*a*,*b*; Case et al., 2016; Ellison et al., 2019). Specifically, Ellison et al. (2019) proposed two statistical criteria for candidate foundation tree species based on their size-frequency and abundance-diameter distributions, and on their spatial effects of on the alpha diversity (as Hill numbers: Chao et al., 2014) and beta diversity (e.g., Bray-Curtis dissimilarity) of co-occurring species.

The first criterion is that candidate foundation tree species are outliers from the expected “reverse-J” size-frequency distribution observed in virtually all assemblages of co-occurring species (Loehle, 2006). The departure from expected size-frequency relationships reflects the abundance of foundation species and their relatively large sizes that lead to their disproportionate influence on overall community structure. We refer to this criterion as the “outlier criterion”.

The second criterion (the “diversity criterion”) is that the size or abundance of candidate foundation tree species should be negatively associated with species diversity (alpha diversity) of other woody plants at local (small) spatial scales but positively associated with species turnover (beta diversity) across large forest plots or stands (Ellison et al., 2019). The negative spatial association between the size or abundance of foundation tree species with local diversity of co-occurring woody species results simply from the foundation species occupying most of the available space in a standard 20*×*20-m forest plot (or, in fact, any relatively small plot). In contrast, the positive spatial association between the size or abundance of a foundation tree species with beta diversity results from it creating patchy assemblages at landscape scales. For example, forest stands dominated by foundation species such as *Tsuga canadensis* in eastern North America or *Pseudotsuga menziesii* in western North America manifest themselves as distinctive patches on the landscape. When these foundation species decline or are selectively harvested, the landscape is homogenized and beta diversity declines. Indeed, Ellison et al. (2019) suggested that the preservation of landscape diversity may be the most important reason to protect and manage foundation tree species before they decline or disappear.

We emphasize that the application of these criteria to identify candidate foundation species leads to the hypothesis that a particular taxon may be a foundation species, not that it is one. Asserting that a species is a foundation species requires additional observational and, ideally, experimental evidence (Ellison, 2014, 2019). Indeed, we derived these two statistical criteria after more than a decade of observational and experimental studies of *Tsuga canadensis*-dominated forests in New England, USA that lend strong support for the hypothesis that *T. canadensis* is a foundation species (Orwig et al., 2013; Ellison, 2014). These criteria subsequently were applied to five additional forest dynamics plots in the western hemisphere (Buckley et al., 2016*b*; Ellison et al., 2019) with encouraging results. Here, we apply these criteria to 12 large forest dynamics plots in China that range from cold-temperate boreal forests to tropical rain forests. These plots are all part of the Chinese Forest Biodiversity Monitoring Network (CForBio)^2^, itself a part of the ForestGEO network.

There are two, fundamentally new contributions of this work. First, we test the hypothesis that foundation tree species should be uncommon or absent in subtropical and tropical forests. Empirical support for particular trees having foundational roles in forests is strongest for temperate forests (Schweitzer et al., 2004; Whitham et al., 2006; Ellison, 2014; Tomback et al., 2016) and low-diversity tropical forests (Ellison et al., 2005), and Ellison et al. (2005) hypothesized that foundation tree species would be less likely in species-rich tropical forests (Ellison et al., 2019). Second, the application of our statistical criteria yield new insights into ecological patterns and processes not only for China, but also concerning similarities between the floras of East Asia and Eastern North America (Tiffney, 1985; Pennington et al., 2004).

## Methods

### Forest dynamics plots in China

We used data from 12 of the 17 CForBio plots in our exporlation of candidate foundation species in Chinese forests (Table 1). These plots span ¿26 degrees of latitude and include: the 9-ha broad-leaved Korean pine mixed forest plot at Liangshui in the Xiaoxing’an Mountains of Heilongjiang Province; the 25-ha *Taxus cuspidata*-dominated forest in the Muling Nature Reserve, also in Heilongjiang Province; the 25-ha deciduous broad-leaved Korean pine mixed forest plot on Changbai Mountain in Jinlin Province; the 20-ha warm-temperate deciduous broad-leaved forest plot on Dongling Mountain in Beijing; the 25-ha subtropical evergreen broad-leaved forest plot on Tiantong Mountain in Zhejiang Province; the 25-ha mid-subtropical mountain evergreen and deciduous broad-leaved mixed forest plot on Badagong Mountain in Hunan province; the 24-ha subtropical evergreen broad-leaved forest plot on Gutian Mountain in Zhejiang Province; 20-ha lower subtropical evergreen broad-leaved forest plot on Dinghu Mountain in Guangdong Province; the 25-ha cold-temperate spruce-fir forest plot on Yulong Snow Mountain in Yunnan Province; the 25-ha karst evergreen and deciduous broad-leaved mixed forest plot at Mulun in the Guangxi Zhuang Autonomous Region; the 15-ha karst seasonal rain-forest plot at Nonggang, also in the Guangxi Zhuang Autonomous Region; and the 20-ha tropical forest plot at Xishuangbanna in Yunnan Province.

**Table 1:**
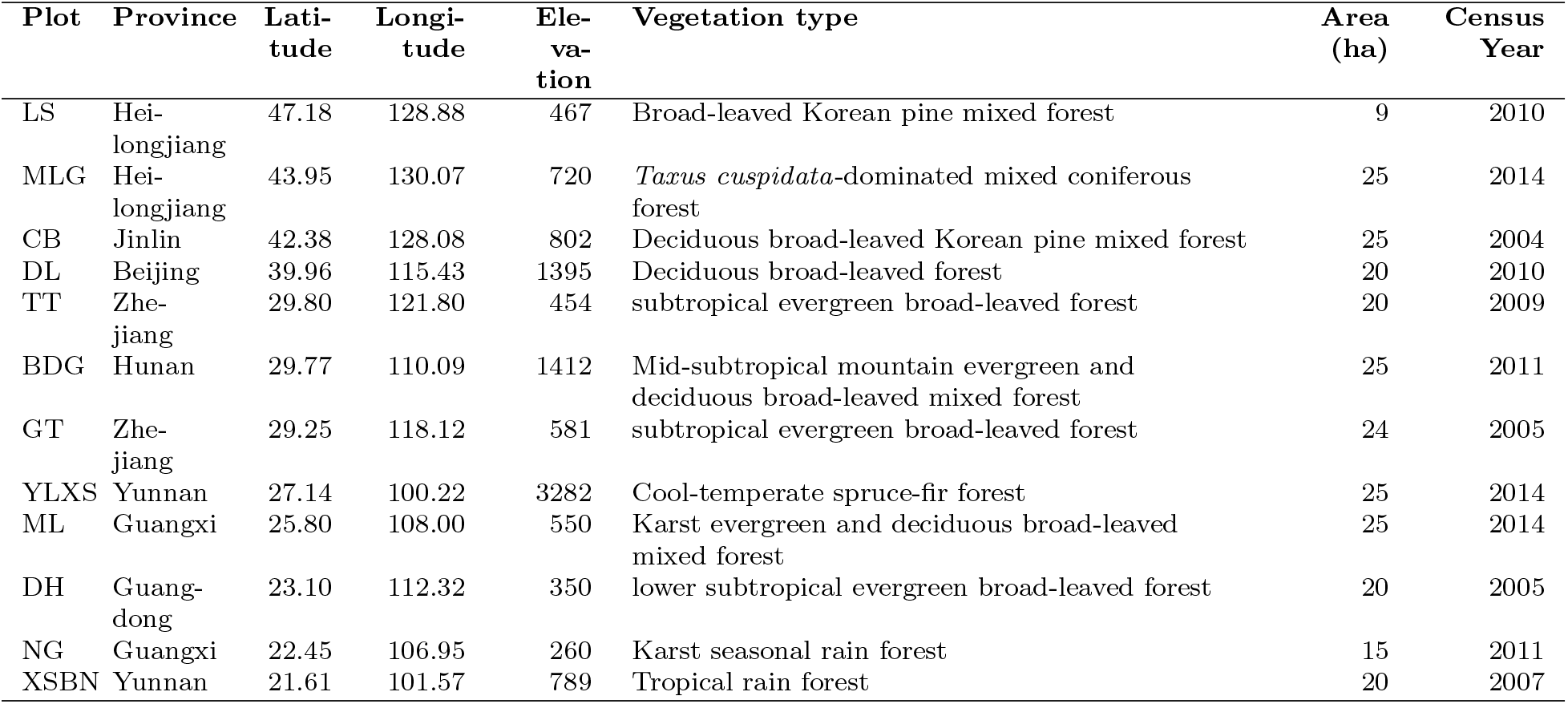
Geographic data for CForBio forest dynamics plots studied here. Latitude and longitude are in *^◦^*N and *^◦^*E, respectively; elevation is in meters above sea level (m a.s.l.); area is in hectares (ha), and census year is the year of the first census of the plot.

| Plot | Province | Latitude | Longitude | Elevation | Vegetation type | Area (ha) | Census Year |
| --- | --- | --- | --- | --- | --- | --- | --- |
| LS | Heilongjiang | 47.18 | 128.88 | 467 | Broad-leaved Korean pine mixed forest | 9 | 2010 |
| MLG | Heilongjiang | 43.95 | 130.07 | 720 | *Taxus cuspidata*-dominated mixed coniferous forest | 25 | 2014 |
| CB | Jinlin | 42.38 | 128.08 | 802 | Deciduous broad-leaved Korean pine mixed forest | 25 | 2004 |
| DL | Beijing | 39.96 | 115.43 | 1395 | Deciduous broad-leaved forest | 20 | 2010 |
| TT | Zhejiang | 29.80 | 121.80 | 454 | subtropical evergreen broad-leaved forest | 20 | 2009 |
| BDG | Hunan | 29.77 | 110.09 | 1412 | Mid-subtropical mountain evergreen and deciduous broad-leaved mixed forest | 25 | 2011 |
| GT | Zhejiang | 29.25 | 118.12 | 581 | subtropical evergreen broad-leaved forest | 24 | 2005 |
| YLXS | Yunnan | 27.14 | 100.22 | 3282 | Cool-temperate spruce-fir forest | 25 | 2014 |
| ML | Guangxi | 25.80 | 108.00 | 550 | Karst evergreen and deciduous broad-leaved mixed forest | 25 | 2014 |
| DH | Guangdong | 23.10 | 112.32 | 350 | lower subtropical evergreen broad-leaved forest | 20 | 2005 |
| NG | Guangxi | 22.45 | 106.95 | 260 | Karst seasonal rain forest | 15 | 2011 |
| XSBN | Yunnan | 21.61 | 101.57 | 789 | Tropical rain forest | 20 | 2007 |

The 9-ha Liangshui plot (“LS”; 47.18 *^◦^*N, 128.88 *^◦^*E) was established in 2005. This plot is located in the Liangshui National Reserve, which has been spared from logging and other major disturbance since 1952 (Liu et al., 2014), and represents the climax vegetation type of Northeast China (Xu and Jin, 2013). It is considered to be one of the most typical and intact mixed broad-leaved-Korean pine forests in China. The plot has an elevational range from 425 to 508 m a.s.l, a mean annual temperature of *−*0.3*^◦^*C, and receives on average 676 mm of precipitation annually. In the first census in 2010, 21,355 individuals stems in 48 species, 34 genera, and 20 families were recorded. The average age of the overstory trees was approximately 200 years (Liu et al., 2014). The “reverse-J” diameter distribution of all individuals in LS suggested that the forest was regenerating well. The dominant tree species at LS is *Pinus koraiensis*. Major associated tree species include *Tilia amurensis, T. mandshurica, Betula costata*, and *Fraxinus mandshurica* (Xu and Jin, 2013).

The 25-ha Muling plot (“MLG”; 43.95 *^◦^*N, 130.07 *^◦^*E) was established in 2014 within the Muling Nature Reserve. The elevation within the plot varies from 658–781 m, the average annual temperature is *−*2*^◦^*C, and the average annual precipitation is 530 mm. Muling is a typical middle-aged, multi-storied, uneven aged forest. Dominant tree species are *Tilia amurensis, Pinus koraiensis, Acer mono, Abies nephrolepis* and *Betula costata*. 63,877 individuals belonging to 22 families, 38 genera, and 57 woody species were recorded at the first census, including the nationally endangered *Taxus cuspidata* (Diao et al., 2016). The average dbh of all woody stems in MLG at the first census was 7.8 cm.

The 25-ha Changbai Mountain plot (“CB”; 42.28 *^◦^*N, 128.08 *^◦^*E), established in 2004, was the first temperate forest dynamics plot in the ForestGEO network. It is considered to be a typical old-growth, multi-storied, uneven-aged forest, and has neither been logged nor suffered other severe human disturbances since 1960 (Wang et al., 2010). The average annual temperature at CB is 3.6 *^◦^*C and average annual precipitation is 700 mm. The terrain of CB is relatively even, with elevations ranging from 791 to 809 m a.s.l. The height of the main canopy species is *≈*30 m, and the oldest trees are *≈*300 years old. In the first census, 38,902 individuals in 52 species representing 32 genera and 18 families were recorded. The most common species at CB are *Pinus koraiensis, Tilia amurensis, Quercus mongolica*, and *Fraxinus mandshurica* (Hao et al., 2008). The most abundant eight species accounted for 83.4% of the total individuals in the plot (Wang et al., 2010).

The 20-ha Dongling Mountain plot (“DL”; 39.96 *^◦^*N, 115.43 *^◦^*E), established in 2010, is in a warm temperate deciduous broad-leaved forest. The average annual temperature at DL is 4.8 *^◦^*C and it receives 500–650 mm of precipitation each year. The mean elevation of the plot is 1395 m, but the terrain is relatively steep with an elevation change of 219 m and slopes ranging from 20–60*^◦^* ^(^Liu et al., 2011). In the first census, 52,316 individuals in 58 species, 33 genera, and 18 families were recorded. The dominant species are all deciduous trees, and include *Quercus wutaishanica, Acer mono*, and *Betula dahurica* (Liu et al., 2011). The most common five species in the plot comprised 61% of all individuals, whilst the most common 20 species comprised 92% of all individuals (Liu et al., 2011).

The 20-ha Tiantong plot (“TT”; 29.80 *^◦^*N, 121.80 *^◦^*E) represents a typical lower subtropical evergreen broad-leaf forest. It was established in 2009 within the core area of the Ningbo Tiantong National Forest Park. Mean annual temperature at TT is 16.2 *^◦^*C and mean annual rainfall is 1375 mm. There have been some typhoon-caused landslides in some parts of the plot (Yang et al., 2011), but it is otherwise considered to be free from human disturbance (Yan et al., 2018). Like Dongling Mountain, TT has a large elevational change across the plot, ranging from 304 to 603 m a.s.l. In the first census, 94,603 individuals in 152 species, 94 genera, and 51 families were recorded. The dominant species are *Eurya loquaiana, Litsea elongata*, and *Choerospondias axiliaris* (Yang et al., 2011).

The 25-ha Badagong Mountain plot (“BDG”; 29.77 *^◦^*N, 110.09 *^◦^*E), established in 2011, is located near the center of distribution of the oak genus *Fagus*. This plot is within the north subtropical mountain humid monsoon climate; the average annual temperature is 11 *^◦^*C and average annual rainfall is 2105 mm (Lu et al., 2013). The dominant trees are a mixture of evergreen (*Cyclobalanopsis multinervis, C. gracilis*, and *Schima parvflora*) and deciduous species (*Fagus lucida, Carpinus fargesii*, and *Sassafras tzumu*). During the first census, 186,556 individuals, belonging to 53 families, 114 genera, and 232 species were recorded (Qin et al., 2018). There were 38 species with ¿1000 individuals, most in the shrub layer (Lu et al., 2013).

The 24-ha Gutian Mountain plot (“GT”; 29.25 *^◦^*N, 118.12 *^◦^*E) was established in 2005 as representing a typical mid-subtropical evergreen broad-leaved forest (Legendre et al., 2009). Like the other montane plots, GT has a broad elevational range (446–715 m a.s.l.) with steep topography (slopes 12–62*^◦^*). Average annual temperature at GT is 15.3 *^◦^*C and average annual rainfall is 1964 mm. In the first census, 140,700 individuals in 159 species, 104 genera, and 49 families were recorded. Dominant species at GT include *Castanopsis eyrei* and *Schima superba* (Legendre et al., 2009).

The 25-ha Yulong Snow Mountain plot (“YLXS”; 27.14 *^◦^*N, 100.22 *^◦^*E), established in 2014, is at the highest elevation (3282 m a.s.l.) of the 12 plots we studied. Although the latitude of this plot is very low, the climate of this coniferous forest plot is cold-temperate because of its high elevation. The average annual temperature at YL is 5.5 *^◦^*C and annual precipitation is 1588 mm (Huang et al., 2017). In the first census, 47,751 individuals in 62 species, 41 genera, and 26 families were recorded, dominated by *Berberis fallax* and *Abies forrestii* (Huang et al., 2017).

The 25-ha Mulun plot (“ML”; 25.80 *^◦^*N, 108.00 *^◦^*E), also established in 2014, is within the Mulun National Natural Reserve. The mean annual temperature at ML is 19.3 *^◦^*C, and the average annual rainfall is 1529 mm. The terrain of the plot is complex and varied. Rock exposure exceeds 60% and soil thickness ¡30 cm in most areas. In the first census, 108,667 individuals in 227 species, 147 genera, and 61 families were recorded (Lan et al., 2016). The dominant species are *Crytocarya microcarpa, Itoa orientalis, Platycarya longipes*, and *Lindera communis* (Lan et al., 2016).

The 20-ha Dinghu Mountain plot (“DH”; 23.10 *^◦^*N, 112.32 *^◦^*E), established in 2005, has an average annual temperature of 20.9 *^◦^*C and average annual precipitation of 1927 mm. This steep, subtropical evergreen forest spans an elevational range of 230–470 m with very steep slopes (30–50*^◦^*). The first census recorded 71,617 individuals in 210 species, 119 genera, and 56 families (Ye et al., 2008). The three canopy-dominant species in the plot are *Castanopsis chinensis, Schima superba* and *Engelhardtia roxburghiana*, whilst the sub-canopy is dominated by *Syzgium rehderianum* and *Craibiodendron scleranthum* var. *kwangtungense* (Ye et al., 2008).

The 15-ha Nonggang plot (“NG”; 22.45 *^◦^*N, 106.95 *^◦^*E), established in 2011, is in a hot-spot of biodiversity in China. This region is characterized by highly vulnerable and spectacular limestone karst systems. Average annual temperature at NG is 21.5 *^◦^*C and average annual precipitation is 1350 mm. The first census recorded 66,718 individuals in 223 species, 153 genera, and 54 families (Lan et al., 2016). Eight of the recorded species are protected throughout China, 30 are endemic to Guangxi province, and three were new records for China. Representative tree species in NG include *Excentrodendron tonkinense, Cephalomappa sinensis, Deutzianthus tonkinensis*, and *Garcinia paucinervis*.

The 20-ha Xishuangbanna plot (“XSBN”; 21.61 *^◦^*N, 101.57 *^◦^*E), established in 2007, is the southernmost CForBio site and is at the northern limit of typical southeast Asian tropical rain forests. It receives 1532 mm of precipitation annually and has an average annual temperature of 21 *^◦^*C. The tropical seasonal rain forest in XSBN is one of the most species-rich forest ecosystems in China. At the first census, 95,834 individuals in 468 species, 213 genera, and 70 families were recorded (Lan et al., 2008). The canopy height of this forest is 50–60 m. The dominant emergent tree species is *Parashorea chinensis*. Subcanopy layers of the forest are dominated by *Sloanea tomentosa, Pometia pinata*, and *Pittosporopsis kerrii*.

### Tree census and measurement

Standard ForestGEO procedures (Condit, 1995) have been used to collect data across all CForBio plots. All woody stems (free-standing trees, shrubs, and lianas) at least 1 cm in diameter at breast height (“dbh”; 1.3 m above the ground level) were tagged, measured, identified to species, and mapped. In all of the plots, the individuals have been censused every 5 years (initial census years in these 12 plots varied between 2004 and 2014; Table 1); we used the first census data from each plot in our analysis.

### The outlier criterion for identifying candidate foundation species

Following Ellison et al. (2019), our first criterion for selecting candidate foundation tree species in each plot was to determine those species that were “outliers” from the typical “reverse-J” distribution of the size-frequency distribution of mean dbh plotted against the number of individuals. We identified outliers by eye rather than fitting a negative exponential distribution with an arbitrary number of parameters to the 12 different size-frequency distributions. This initial screen revealed 2–14 candidate foundation tree species in each of the 12 forest dynamics plots (Fig. 1). The largest number of candidate species occurred in MLG and the fewest were in YLXS. To screen species more expansively and avoid missing other possible candidate foundation species, we also included in our first cut those ten species with the highest importance values (iv = relative abundance + relative density + relative basal area) in each of the plots. Species that were outliers on the size-frequency plots usually had high importance values, but including the latter did expand our initial pool of candidate species to 10–14 species per plot (Table 2).

**Figure 1:**
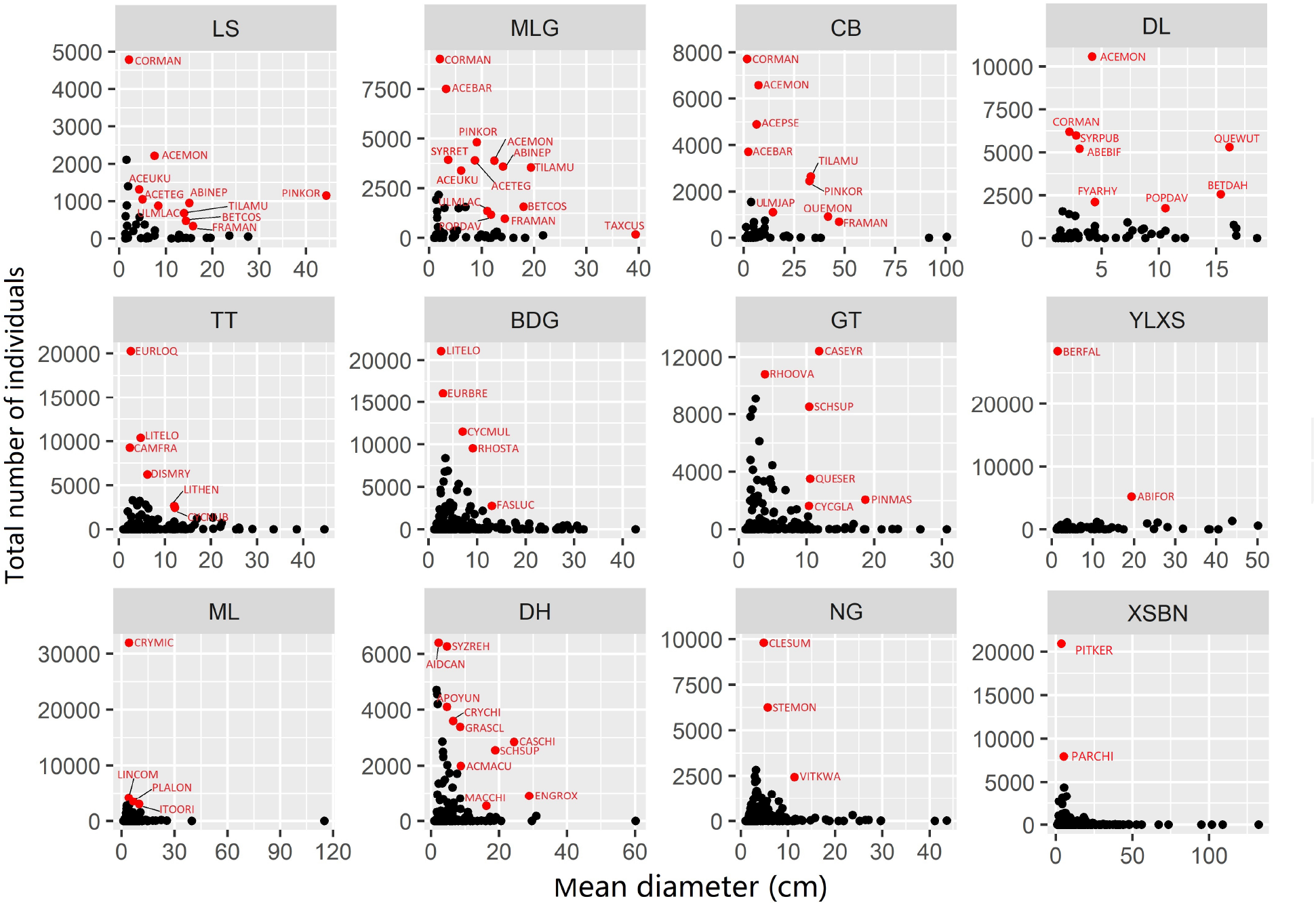
Size (dbh)-frequency distributions of the species in each plot. Species falling outside of the “reverse-J” line (in red) were treated in the first set of candidate foundation species. Plots are ordered left-to-right and top-to-bottom by latitude; species abbreviations are given in Table 2.

**Table 2:**
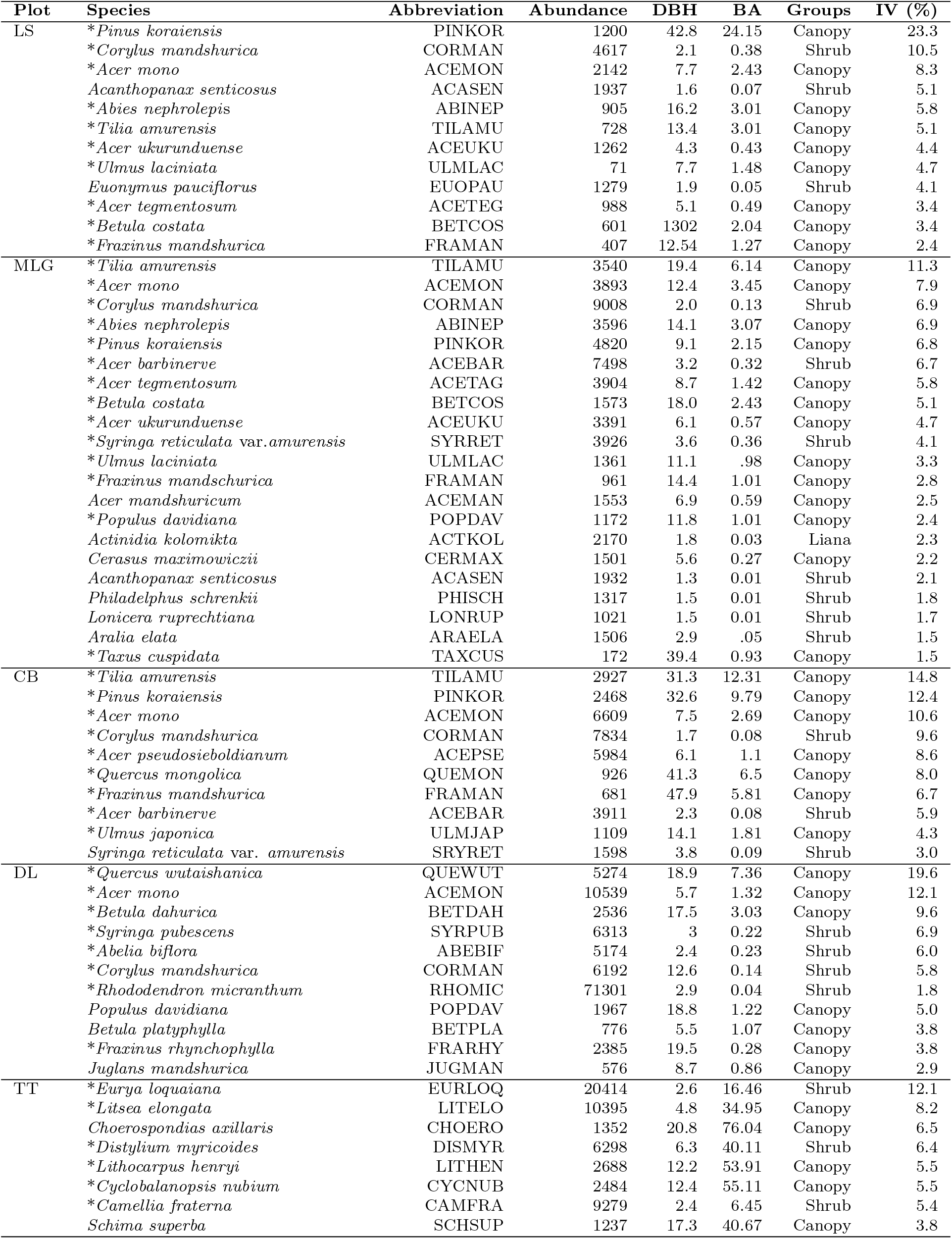
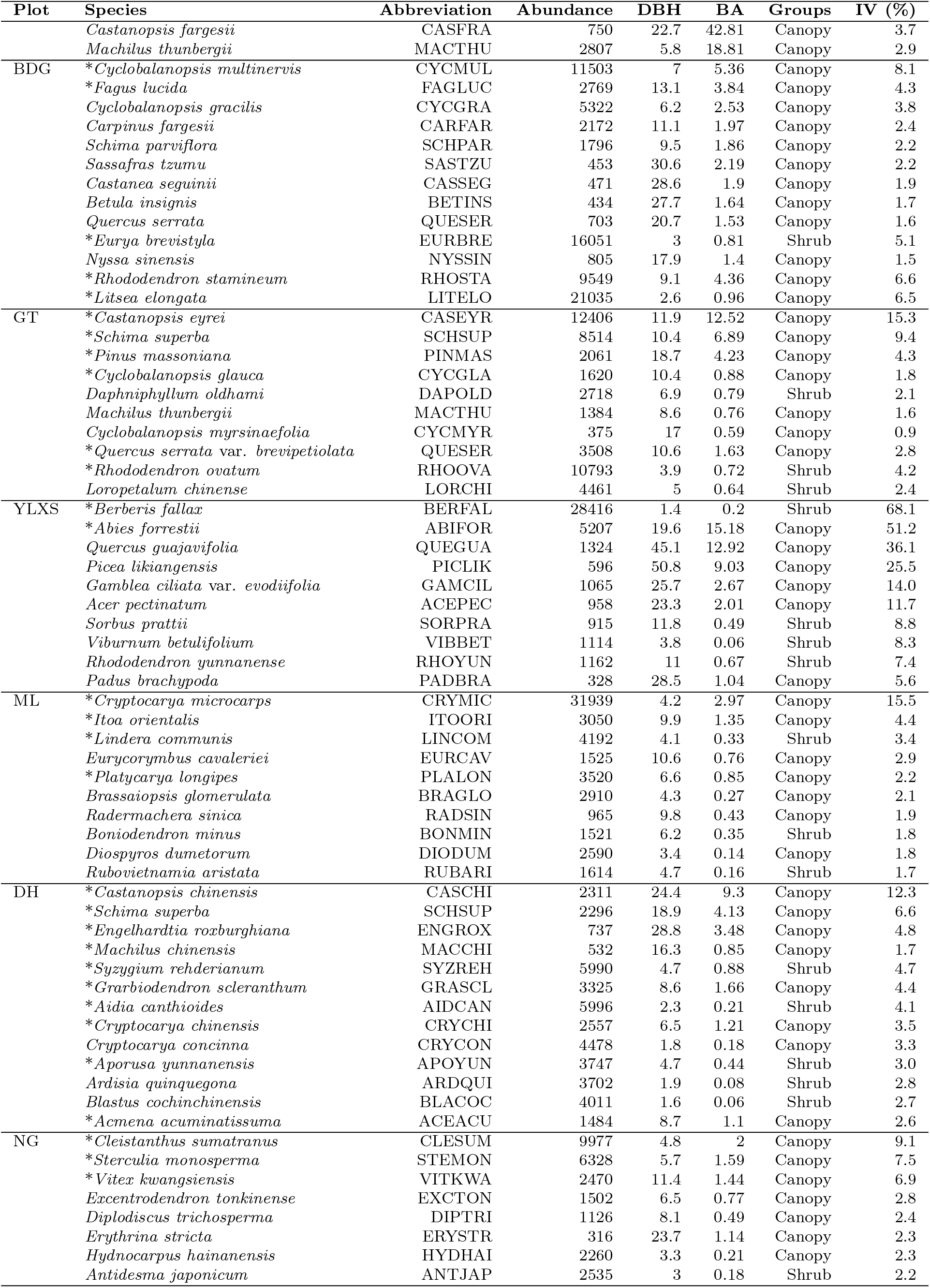
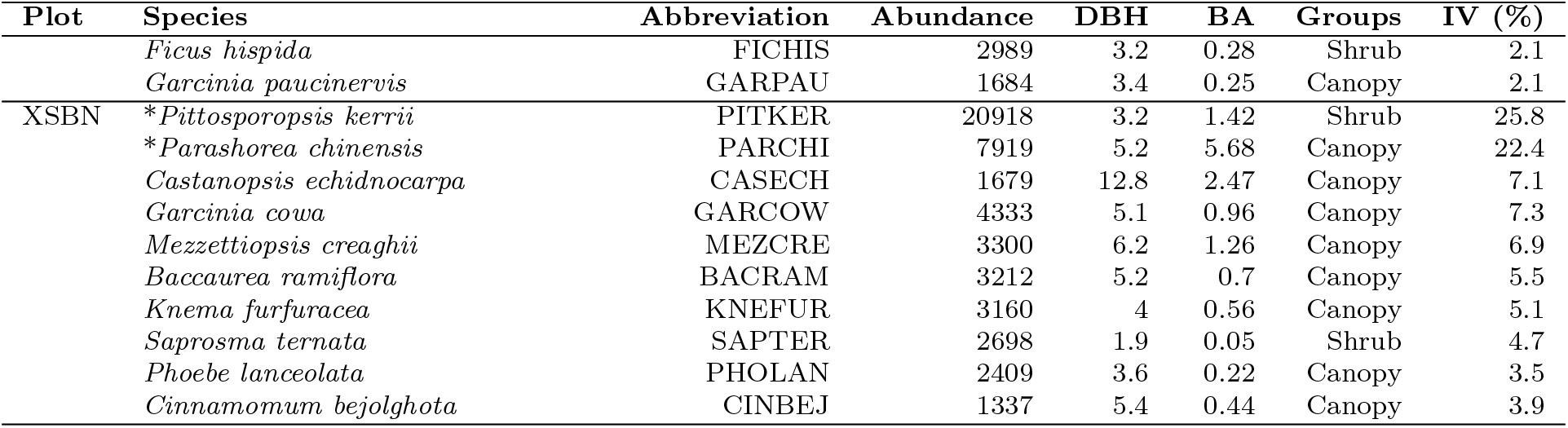
Initial set of candidate foundation species identified as outliers in the abundance-dbh plots (Fig. 1; here marked with an asterisk[*]) and others whose importance values IV) were in the top ten for that plot. Plots are ordered by latitude, and within each plot, species are ordered by IV. Units of diameter (dbh) are cm and units of basal area (BA) are in m^2^/ha.

### The diversity criterion for identifying candidate foundation species

The second, more stringent criterion for identifying candidate foundation species is a negative association between its size (or abundance), and total abundance, three measures of alpha diversity (species richness, Shannon diversity, Inverse Simpson Diversity) of associated woody species *and* a positive association between its size or abundance and beta diversity (Ellison et al., 2019). The three measures of alpha diversity treat all species identically (species richness), down-weight rare species (Shannon diversity), or down-weight common species (inverse Simpson diversity) within subplots. These associations also should be consistent across the plots when calculated at a given spatial grain (*a.k.a.* spatial scale) and at most (ideally all) spatial lags (Buckley et al., 2016*a*; Ellison et al., 2019).

#### Forest structure and species diversity indices

For each plot, we calculated the total basal area, mean basal area, and total number of individuals of each of the candidate foundation tree and shrub species (Table 3) within contiguous 5 *×* 5, 10 *×* 10, and 20 *×* 20-m subplots. For species other than the candidate foundation species, we calculated their total abundance, species richness, Shannon and inverse Simpson diversity indices (as Hill numbers: Chao et al., 2014) and mean Bray-Curtis dissimilarity (overall methods as in Ellison et al., 2019). In all the analysis, we used only the main stem of each individuals (i.e., smaller stems of multi-stemmed individuals were excluded from the analyses). The diversity() and vegdist() functions in the vegan package (Oksanen et al., 2018) of the R software system (R Core Team, 2019) were used for calculating each diversity metric.

**Table 3:** A winnowed list of candidate foundation tree and shrub species (the latter indicated by a plus sign [^+^]) at three different spatial grains (i.e., subplot size) in 12 Chinese forest dynamics plots. Plots are ordered by latitude, and within each plot, candidate foundation species are ordered alphabetically. The two *Acer* species in **bold type** satisfied all aspects of both the outlier and the diversity criteria for candidate foundation species at the given spatial grain. The starred (*) species satisfied the outlier criterion (Fig. 1) and partially satisfied the diversity criterion at the given spatial grain: a positive spatial relationship between candidate foundation species size and beta diversity, and a negative spatial relationship between candidate foundation species size and at least one measure of alpha diversity. The remaining species did not satisfy the outlier criterion but did meet some aspects of the diversity criterion. No species met either foundation species criterion in the BDGS, TTS and YLXS plots at any spatial grain.

| Plot | 5 m | Spatial grain<br>10 m | 20 m |
| --- | --- | --- | --- |
| LS | * *Acer ukurunduense*<br>* *Corylus mandshurica* <sup>+</sup><br>* *Fraxinus mandshurica* | * *Acer ukurunduense*<br>—<br>— | —<br>—<br>— |
| MLG | * ***Acer barbinerve*** <sup>+</sup><br>* *Acer tegmentosum*<br>* ***Acer ukurunduense***<br>* *Corylus mandshurica* <sup>+</sup><br>—<br>* *Taxus cuspidata*<br>* *Tilia amurensis* | * *Acer barbinerve* <sup>+</sup><br>—<br>—<br>—<br>* *Pinus koraiensis*<br>—<br>* *Tilia amurensis* | —<br>—<br>—<br>—<br>* *Pinus koraiensis*<br>—<br>* *Tilia amurensis* |
| CB | * *Acer barbinerve* <sup>+</sup><br>* *Acer pseudosieboldianum*<br>* *Corylus mandshurica* <sup>+</sup><br>*Syringa reticulata* var. *amurensis* <sup>+</sup> | —<br>* *Acer pseudosieboldianum*<br>* *Corylus mandshurica* <sup>+</sup><br>*Syringa reticulata* var. *amurensis* <sup>+</sup> | —<br>—<br>—<br>— |
| DL | *Juglans mandshurica* | — | — |
| TT | — | — | — |
| BDG | — | — | — |
| GT | *Machilus thunbergii*<br>—<br>— | —<br>—<br>— | —<br>* *Pinus massoniana*<br>* *Quercus serrata* var. *brevipetiolata* |
| YLXS | — | — | — |
| ML | *Brassaiopsis glomerulata* | *Brassaiopsis glomerulata* | — |
| DH | * *Aporosa yunnanensis* <sup>+</sup> | * *Aporosa yunnanensis* <sup>+</sup> | * *Aporosa yunnanensis* <sup>+</sup> |
| NG | *Ficus hispida*<sup>+</sup> | *Ficus hispida*<sup>+</sup> | — |
| XSBN | *Mezzettiopsis creaghii* | *Mezzettiopsis creaghii* | *Mezzettiopsis creaghii* |

#### Codispersion analysis

Codispersion describes anisotropic spatial patterns (i.e., different expected values when measured in different directions) of co-occurring variables for given spatial lags and directions (Cuevas et al., 2013). The codispersion coefficient ranges from *−*1 to 1, with positive values indicating a positive spatial association and negative values indicating a negative spatial association for a given spatial lag and direction. These values can be visualized with a codisperison graph (Vallejos et al., 2015). Buckley et al. (2016*a*) introduced codispersion analysis to ecologists through an exploration of spatial patterns of species co-occurrence. That paper also provides a basic introduction to the mathematics of codispersion analysis and codispersion graphs. Buckley et al. (2016*b*) used codispersion analysis to examine spatial patterns of relationships between environmental characteristics and known or candidate foundation tree species. Ellison et al. (2019) used codispersion analysis to quantify spatial effects of candidate foundation tree species on different measures of diversity of associated woody species in six forest dynamics plots in the Americas.

Although we computed codisperison patterns using mean basal area, total basal area, and total abundance of candidate foundation species, we focus our presentation on the codispersion between the total basal area of the candidate foundation species and associated woody plant diversity in the differently-sized (5 *×* 5, 10 *×* 10, and 20 *×* 20-m subplots) contiguous subplots in each of the 12 forest dynamics plots (Ellison et al., 2019); qualitatively similar patterns were observed when using mean basal area or total numbers of individuals of candidate foundation species. For each candidate foundation tree species, we first computed the observed codispersion coefficient between its total basal area and abundance, alpha, and beta diversity of the associated woody species in the subplots. The maximum spatial lag examined for each plot ranged from the length of the subplot to one-fourth of the length of the shortest side of each forest plot, which ensured adequate sample sizes for reliable estimation of codispersion coefficients at the largest spatial lag (Buckley et al., 2016*a*).

Statistical significance of the codispersion coefficients was determined using null model analysis (Buckley et al., 2016*b*; Ellison et al., 2019). Codispersion coefficients for all spatial lags and directions were computed for co-occurrence matrices randomized using a toroidal-shift null model, which maintains the autocorrelation structure of the species and spatial patterns caused by underlying environmental gradients while shifting the associated woody species in random directions and distances (Buckley et al., 2016*b*; Ellison et al., 2019). For each candidate foundation species in each plot, we ran 199 randomizations; significance was determined based on empirical 95% confidence bounds. Calculation of codispersion coefficients and all randomizations were done using custom C and R code written by Ronny Vallejos and Hannah Buckley, respectively.

### Data and code availability

Each of the CForBio plots were established at different times and are scheduled to be (or already have been) censused every five years. To maximize comparability among datasets, we used data collected at the first census for each plot (Table 1). All datasets are available from the ForestGEO data portal https://ctfs.si.edu/datarequest). R code for all analyses is available on GitHub (https://github.com/buckleyhannah/FS_diversity.

## Results

### Candidate foundation species in the CForBio plots

Only two species—the shrub *Acer barbinerve* (Fig. 2, 3) and the congeneric tree *Acer ukurunduense* (Fig. 4, 5)—in one plot—MLG—satisfied both the outlier *and* diversity criteria for all diversity measures for candidate foundation species (Figs. 2–5). For these two species in MLG, both criteria were met only at the 5-m spatial grain (Table 3).

**Figure 2:**
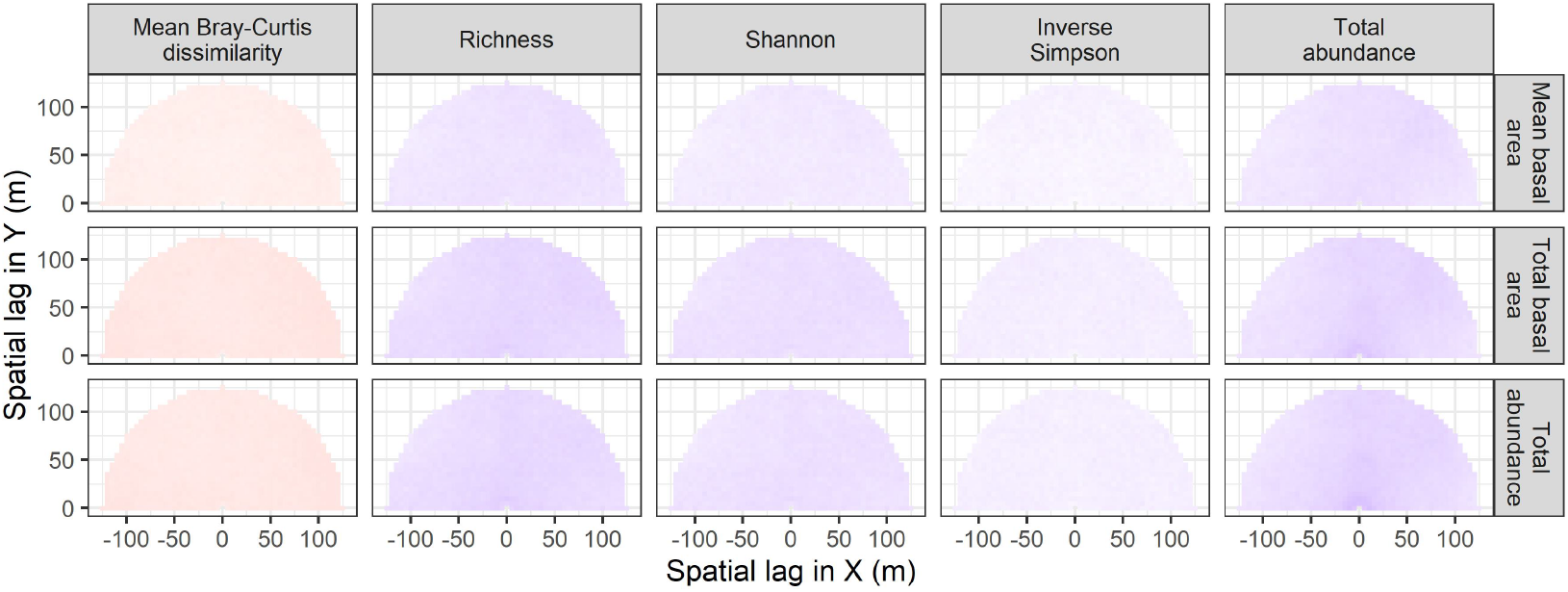
Codispersion between mean basal area, total basal area, or total abundance of *Acer barbinerve* and five different measures of diversity of associated woody species in 5-m subplots in the 25-ha Muling (MLG) plot. Codispersion coefficients were calculated for spatial lags ranging from 0–125 m at 5-m intervals. The values of the codispersion can range from −1 (dark blue) through 0 (white) to 1 (dark red). Statistical significance for codispersion coefficients computed at each spatial lag is shown in Fig. 3.

**Figure 3:**
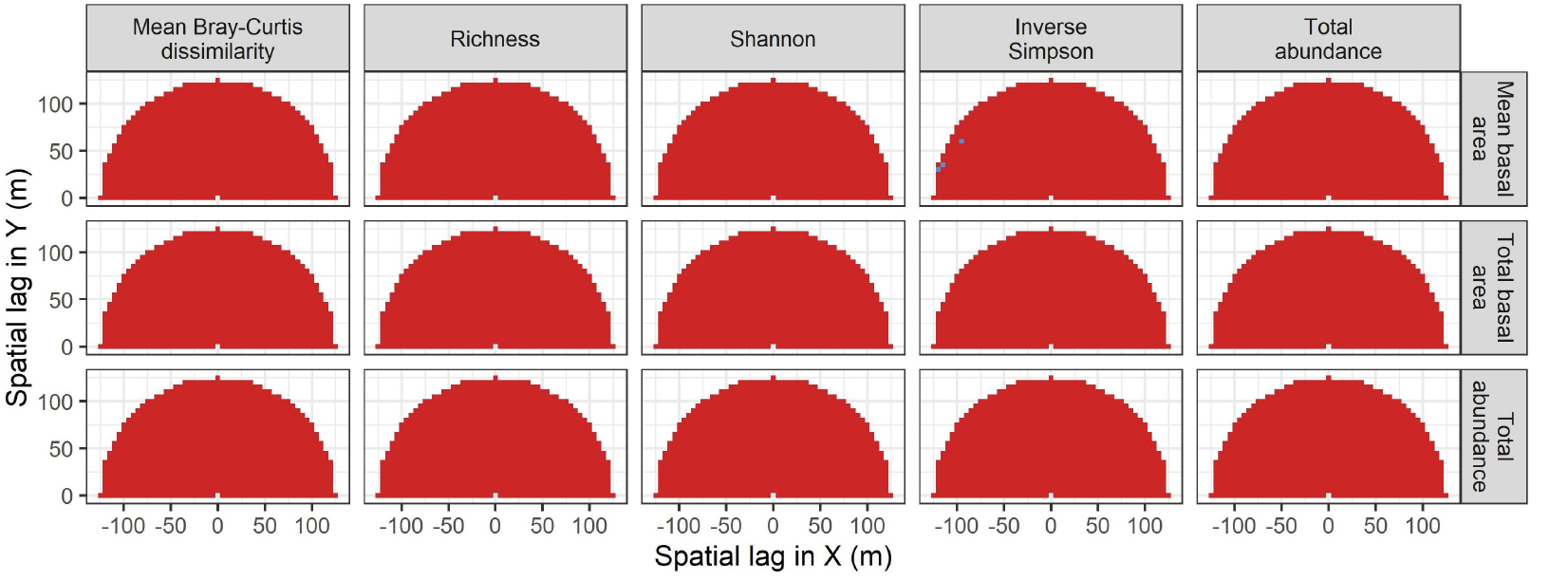
Statistical significance of the codispersion coefficients calculated between basal area or abundance of *Acer barbinerve* and five different measures of diversity of associated woody species in 5-m subplots in the 25-ha Muling (MLG) plot. Statistical significance was determined by comparing observed codispersion at each spatial lag with the distribution of 199 spatial randomizations of a toroidal-shift null model. Red: *P ≤* 0.05; Blue: *P >* 0.05.

**Figure 4:**
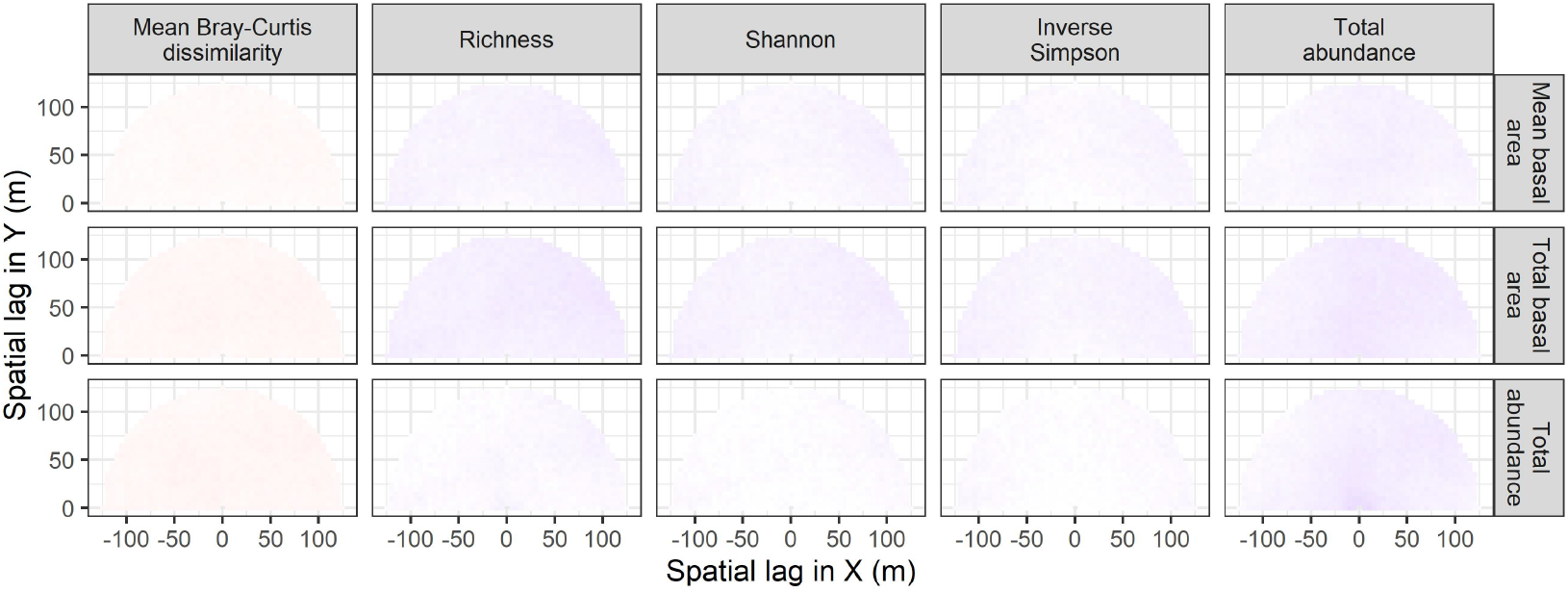
Codispersion between mean basal area, total basal area, or total abundance of *Acer ukurunduense* and five different measures of diversity of associated woody species in 5-m subplots in the 25-ha Muling (MLG) plot. Codispersion coefficients were calculated for spatial lags ranging from 0–125 m at 5-m intervals. The values of the codispersion can range from −1 (dark blue) through 0 (white) to 1 (dark red). Statistical significance for codispersion coefficients computed at each spatial lag is shown in Fig. 5.

**Figure 5:**
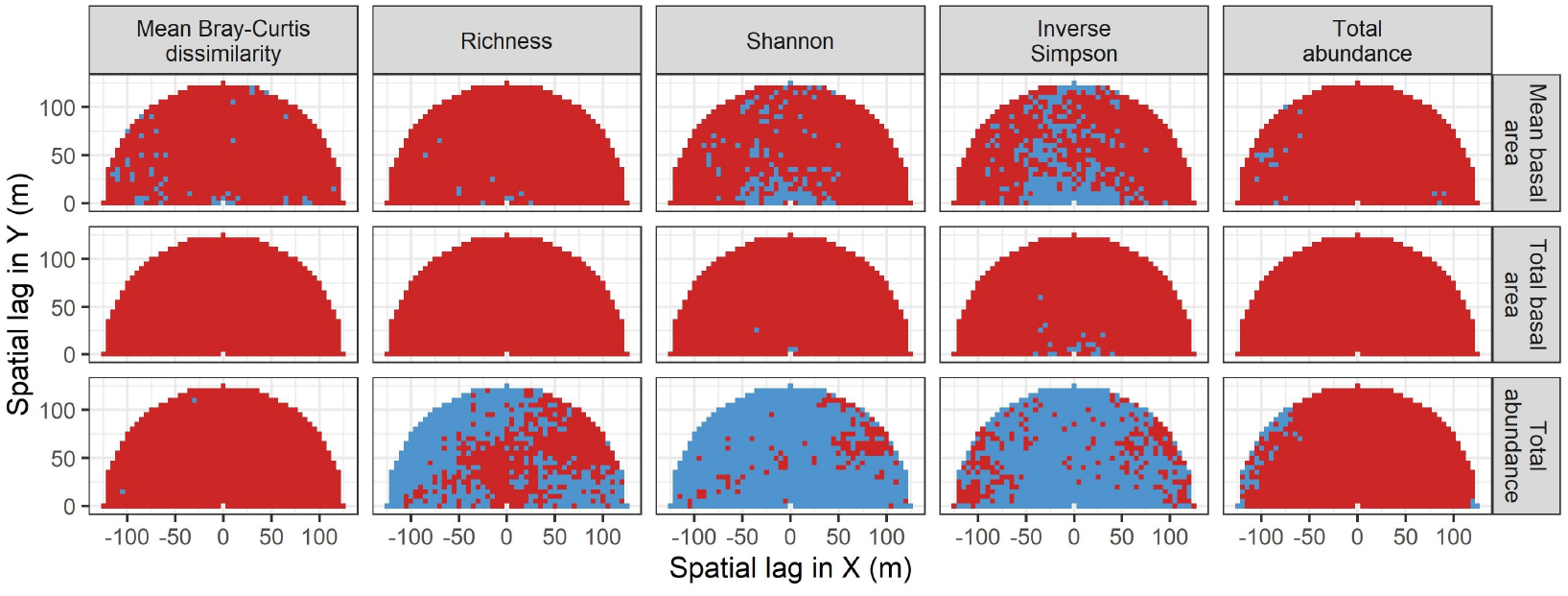
Statistical significance of the codispersion coefficients calculated between basal area or abundance of *Acer ukurunduense* and five different measures of diversity of associated woody species in 5-m subplots in the 25-ha Muling (MLG) plot. Statistical significance was determined by comparing observed codispersion at each spatial lag with a distribution of 199 spatial randomizations of a toroidal-shift null model. Red: *P ≤* 0.05; Blue: *P >* 0.05.

More species were considered as candidate foundation species when we retained the outlier criterion (Fig. 1) but relaxed the diversity criterion to require only a positive spatial relationship between the size of the candidate foundation species and beta diversity *and* a negative spatial relationship between the size of the candidate foundation species and at least one of the alpha-diversity measures (species indicated with an asterisk [*] in Table 3). These additional candidate foundation species included two additional *Acer* species, tree species in the genera *Pinus, Taxus, Fraxinus, Quercus*, and *Tilia*, and two shrubs (*Corylus mandshurica* and *Aporusa yunnanensis*). However, whether we applied the stringent or relaxed diversity criterion, all but three of the candidate foundation species occurred in plots with cool- or cold-temperate climates. The exceptions were the trees *Pinus massoniana* and *Quercus serrata* var. *brevipetiolata* at GT and *Aporusa yunnanensis* at DH; all three of these species occurred in the subtropical evergreen broad-leaved forest plots.

A few of our initial candidate species that had high importance values but were not outliers from the expected size-frequency distributions (unstarred species in Table 2) did partially meet the diversity criterion in both temperate and tropical plots (Table 3). These included *Syringa reticulata* var. *amurensis* at CB, *Juglans mandshurica* at DL, *Machilus thunbergii* at GT, *Brassaiopsis glomerulata* at ML, *Ficus hispida* at NG, and *Mezzettiopsis creaghii* at XSBN.

### Scale-dependence of candidate foundation species

More candidate foundation species—including all species that met at least one of the two criteria—were identified at smaller spatial grains: 15 species at the 5-m grain, 11 at the 10-m grain, and six at the 20-m grain (Table 3). This pattern applied both among and within the plots. Average codispersion between total basal area of the candidate foundation species and Bray-Curtis dissimilarity increased significantly with spatial grain (Fig. 6; raw data in Table 4). In contrast, average codispersion between total basal area of the candidate foundation species and measures of alpha diversity, while generally negative, were more variable and not scale-dependent (Fig. 6; raw data in Table 4).

**Figure 6:**
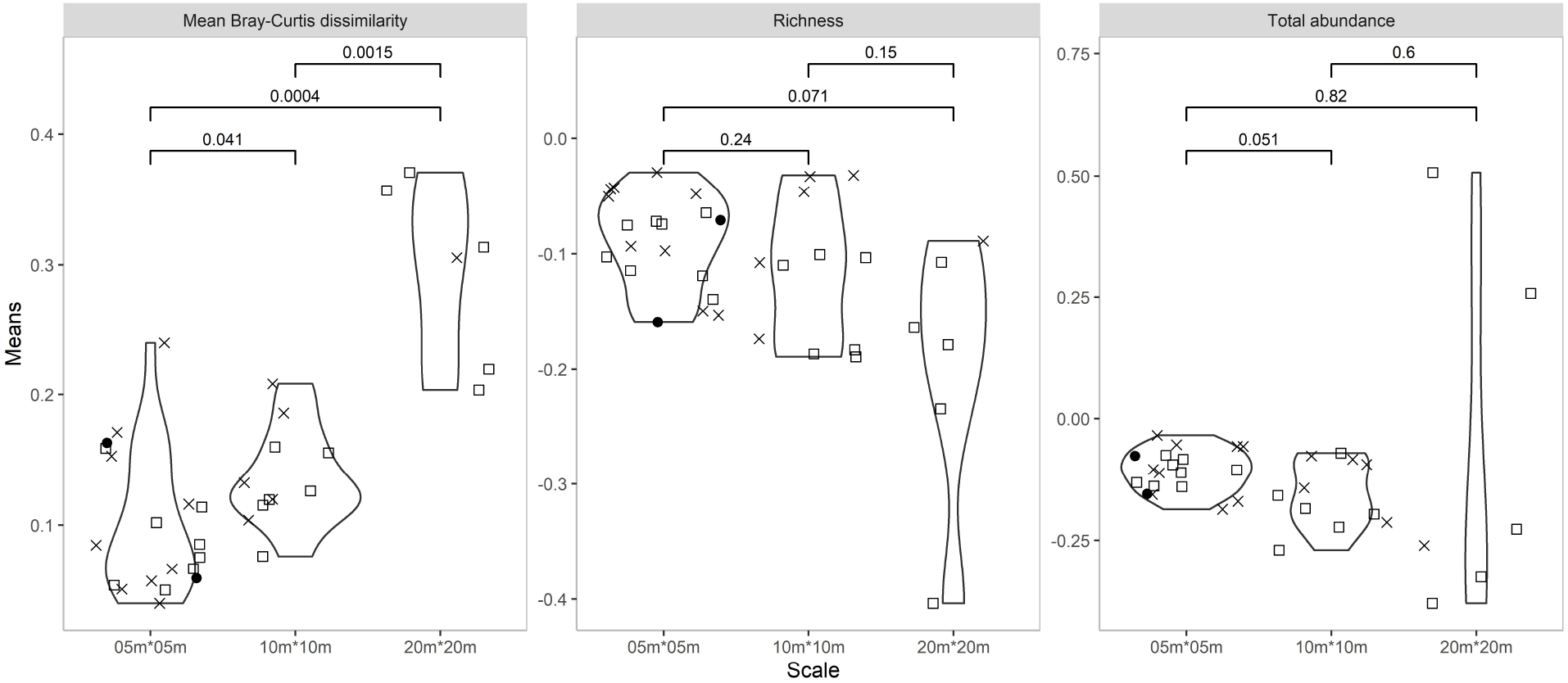
Distribution of average codispersion observed between total basal area of candidate foundation species and Bray-Curtis dissimilarity, species richness, and total abundance of associated woody plant species in continguous 5 × 5-, 10 × 10-, and 20 × 20-m subplots in the twelve CForBio plots. Points indicate mean codispersion values for each candidate foundation species listed in Table 2; solid points indicate the two candidate foundation species in the genus *Acer* that met both the outlier *and* diversity criterion for all indices; hollow squares indicate candidate species that met the outlier criterion and the relaxed diversity criterion; and crosses indicate the remaining candidate foundation species that met only the relaxed diversity criterion. *P* values for comparisons between groups are shown at the top of each panel.

**Table 4:**
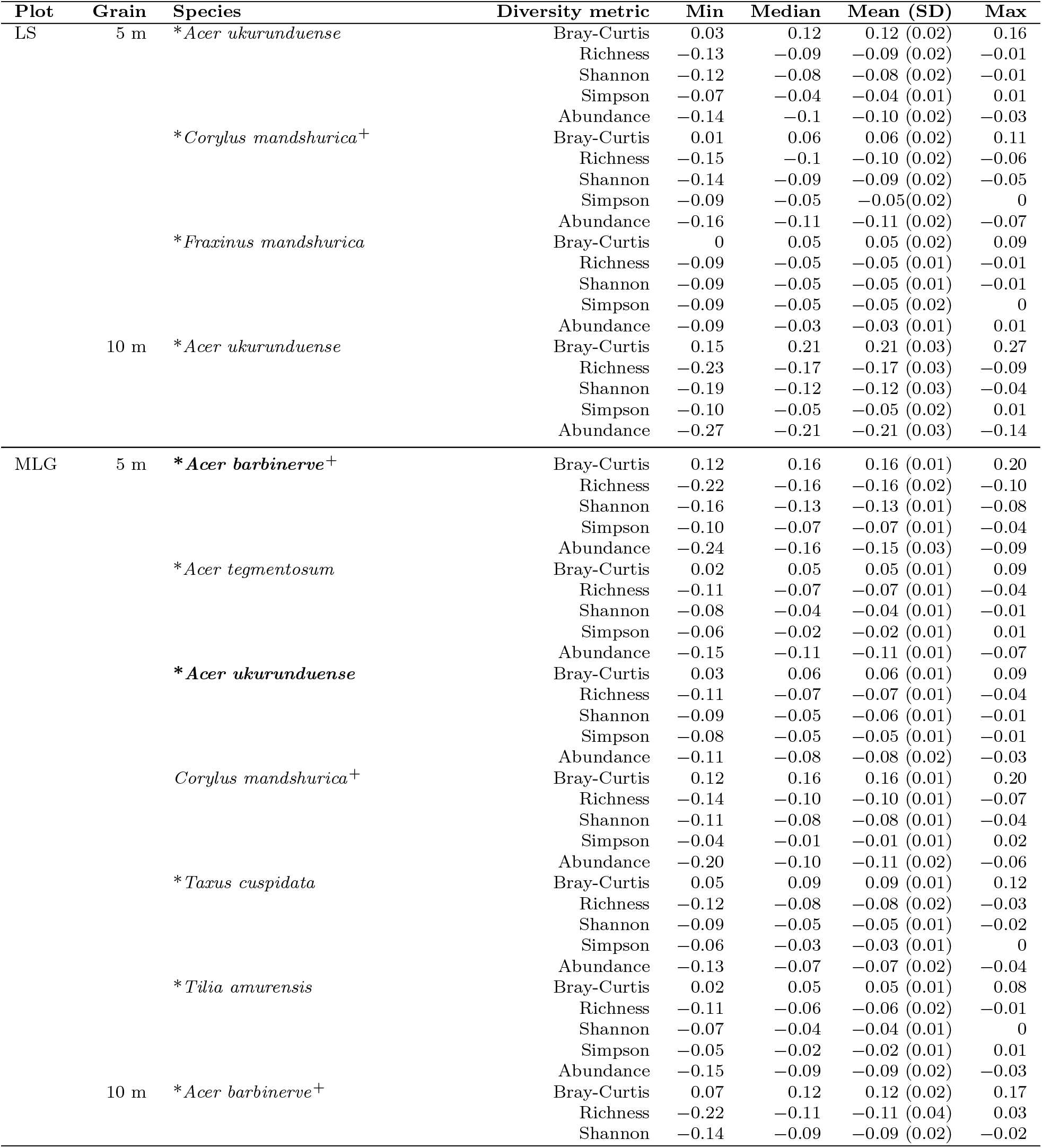
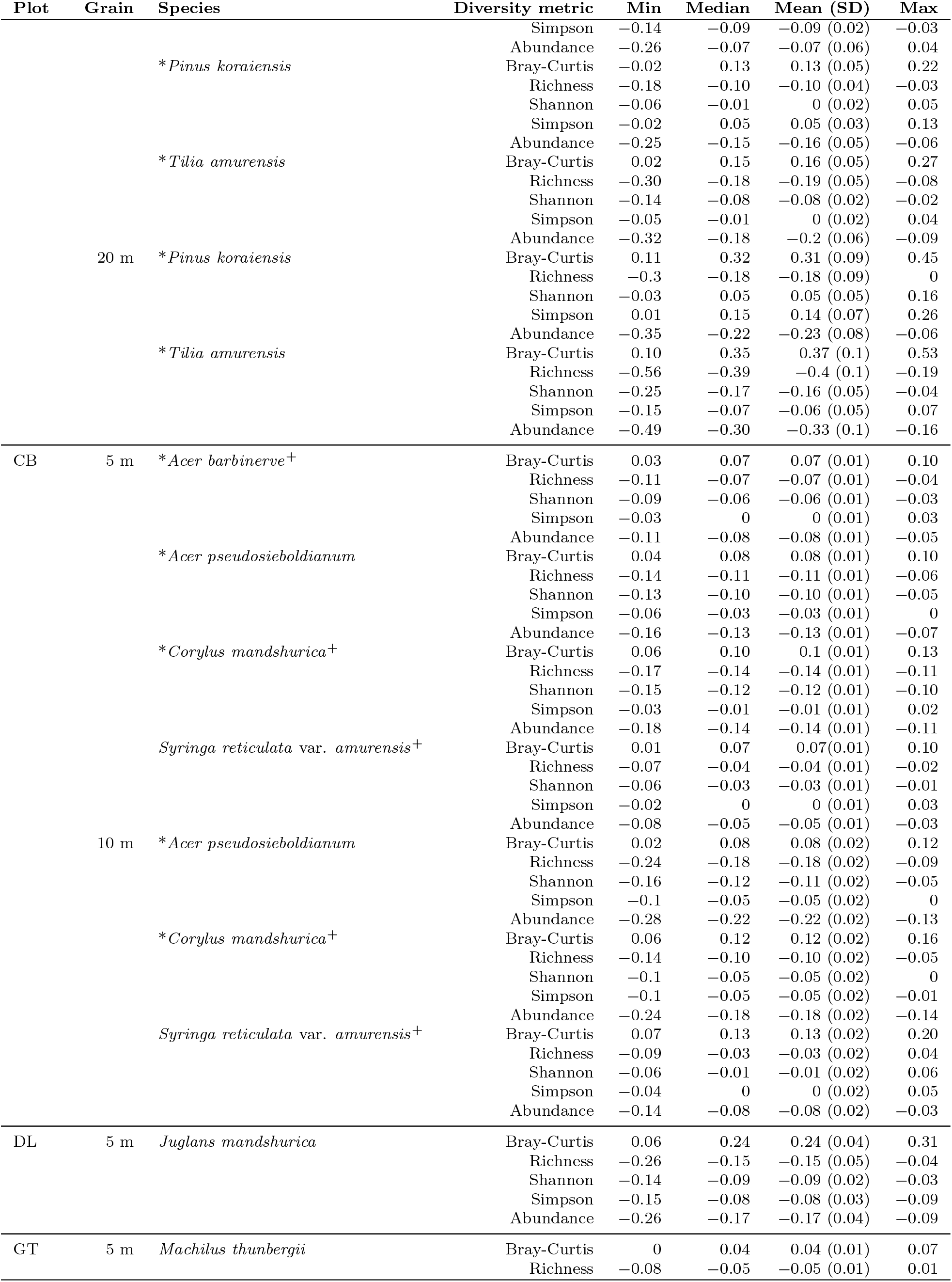
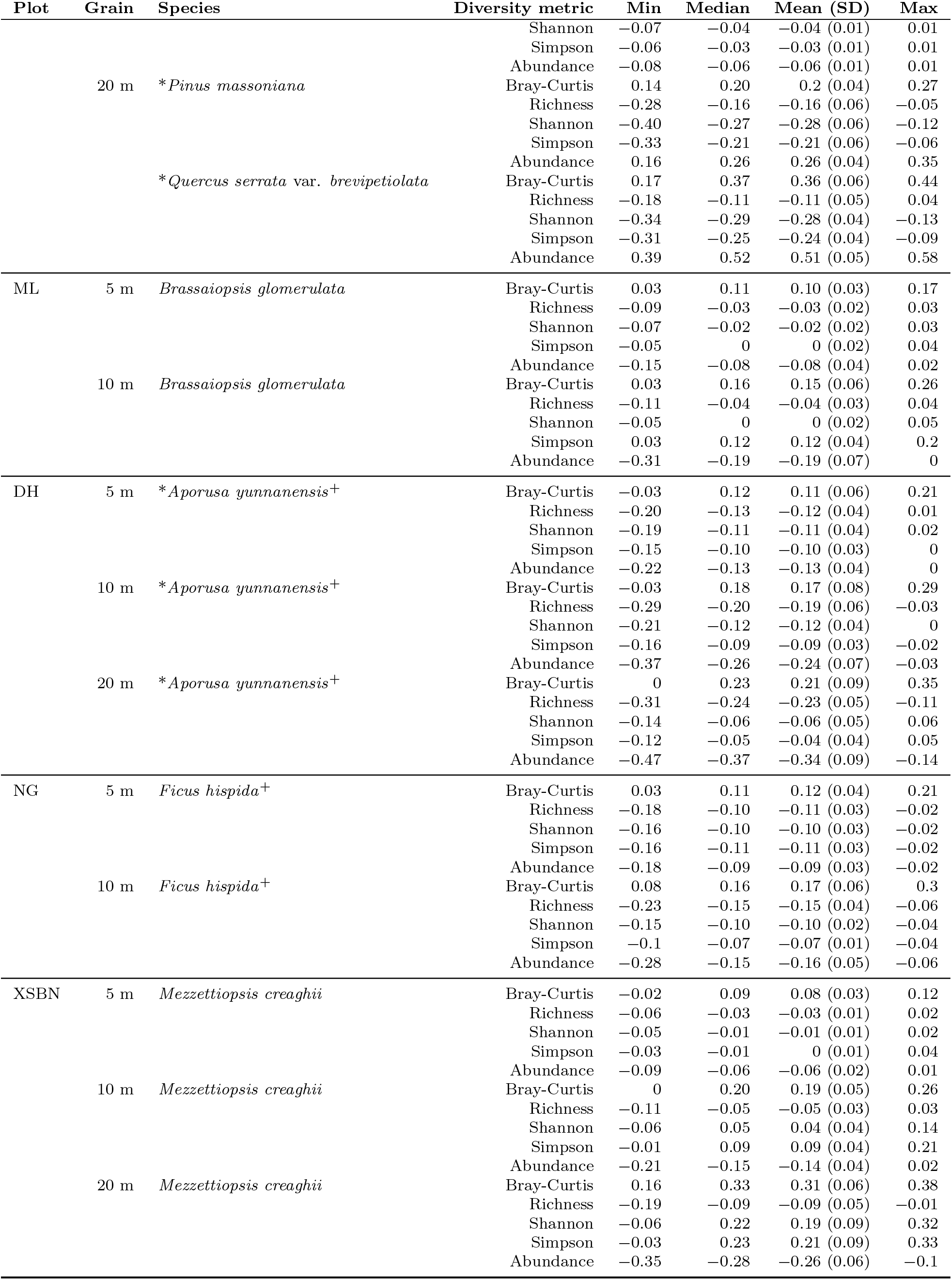
Codisperison statistics for the candidate foundation tree or understory species (the latter indicated by a [^+^]) in each plot at the spatial grain (**Grain**) at which they were identified (species listed in Table 3). As in Table 3, the two *Acer* species in **bold type** satisfied all aspects of both the outlier and the diversity criteria for candidate foundation species at the given spatial grain. The starred (*) species satisfied the outlier criterion (Fig. 1) and partially satisfied the diversity criterion at the given spatial grain: a positive spatial relationship between candidate foundation species size and beta diversity, and a negative spatial relationship between candidate foundation species size and at least one measure of alpha diversity. The remaining species did not satisfy the outlier criterion but did meet some aspects of the diversity criterion. No species met either foundation species criterion in the BDGS, TTS and YLXS plots at any spatial grain. Plots are ordered by latitude, and within each plot, species are grouped alphatically within increasing grain (subplot) sizes. Values are the minimum (**Min**), median (**Median**), mean (**Mean**), one standard deviation of the mean (**SD**), and maximum (**Max**), computed over all spatial lags, of the codispersion between the basal area of the candidate foundation species and all other woody species in square subplots with the length of a side = the spatial grain.

### Candidate foundation species across a latitudinal gradient

Both the number of woody species in each plot that were outliers from the expected size-frequency distribution and the number of candidate foundation species increased with increasing latitude (Fig. 7A, C; slopes = 0.3 and 0.15 species/degree of latitude, respectively; *P* ¡0.01). As expected, within-plot species richness declined significantly with latitude (slope = *−*10.2 species/degree of latitude, *P* ¡0.01), but this relationship was unrelated to the latitudinal pattern in either the number of outliers or the number of candidate foundation species. There were no significant relationships between either the number of outliers or the number of candidate foundation species and within-plot species richness (Fig. 7B, D; *P* = 0.08 and 0.18 respectively).

**Figure 7:**
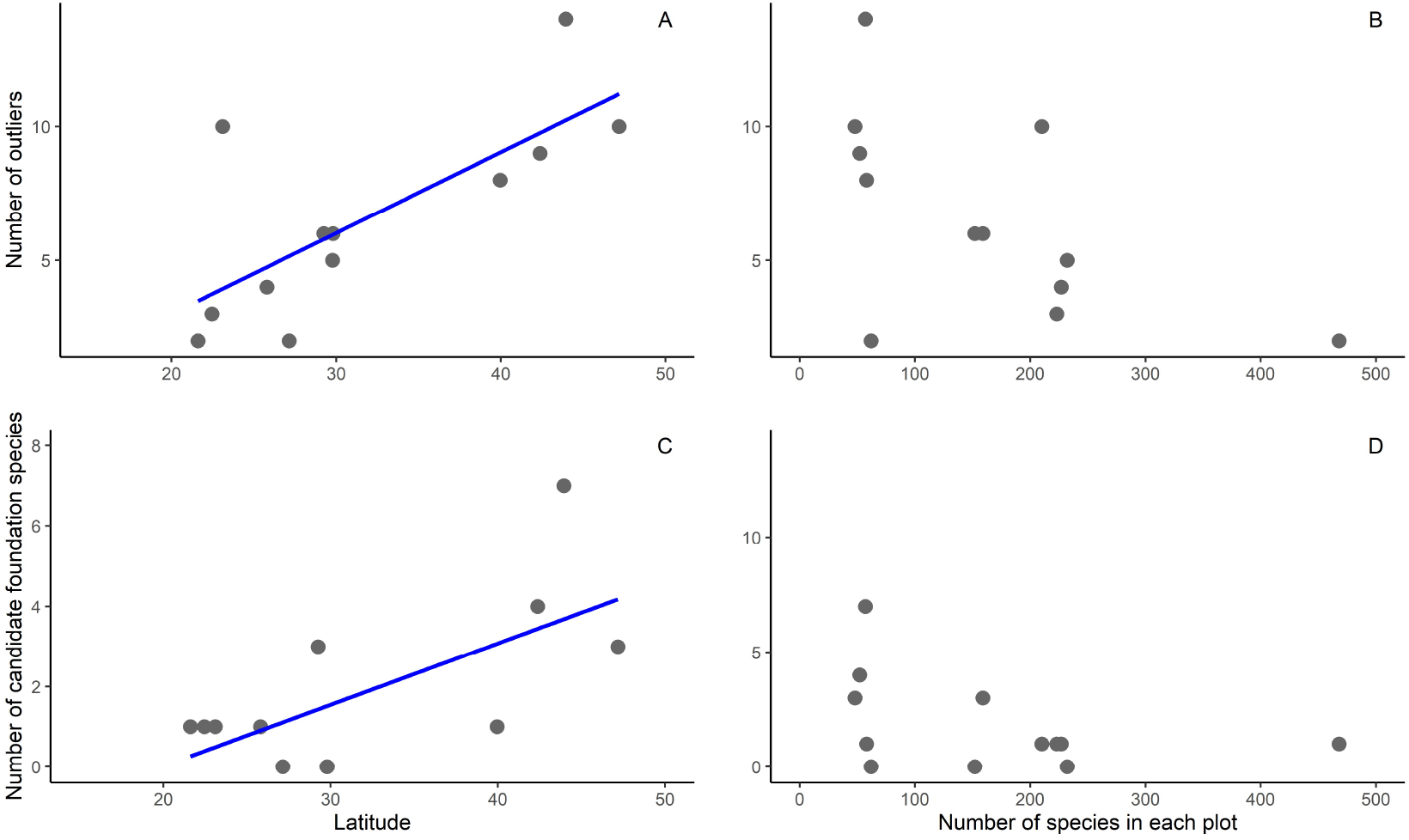
Number of outliers from the expected size-frequency distribution (Fig. 1) and number of candidate foundation species (Table 3) as a function of latitude (**A, C**) or plot-level species richness (**B, D**). See main text for regression statistics.

Spatial association (expressed as codispersion) within each plot between candidate foundation species and total abundance, mean alpha diversities, and mean beta diversity of associated woody species on average did not vary with latitude at any spatial grain (Fig. 8; raw data in Table 4). Quantile regression (to account for potential extreme effects of foundation species) yielded similar results. There were no observed latitudinal patterns in effects of candidate foundation species except for a slight strengthening of the negative effect of candidate foundation species on associated woody species richness and total abundance at the 5-m grain (Fig. 8, *P* = 0.04). Similar results were obtained when understory shrubs were excluded from the analysis (Fig. 9).

**Figure 8:**
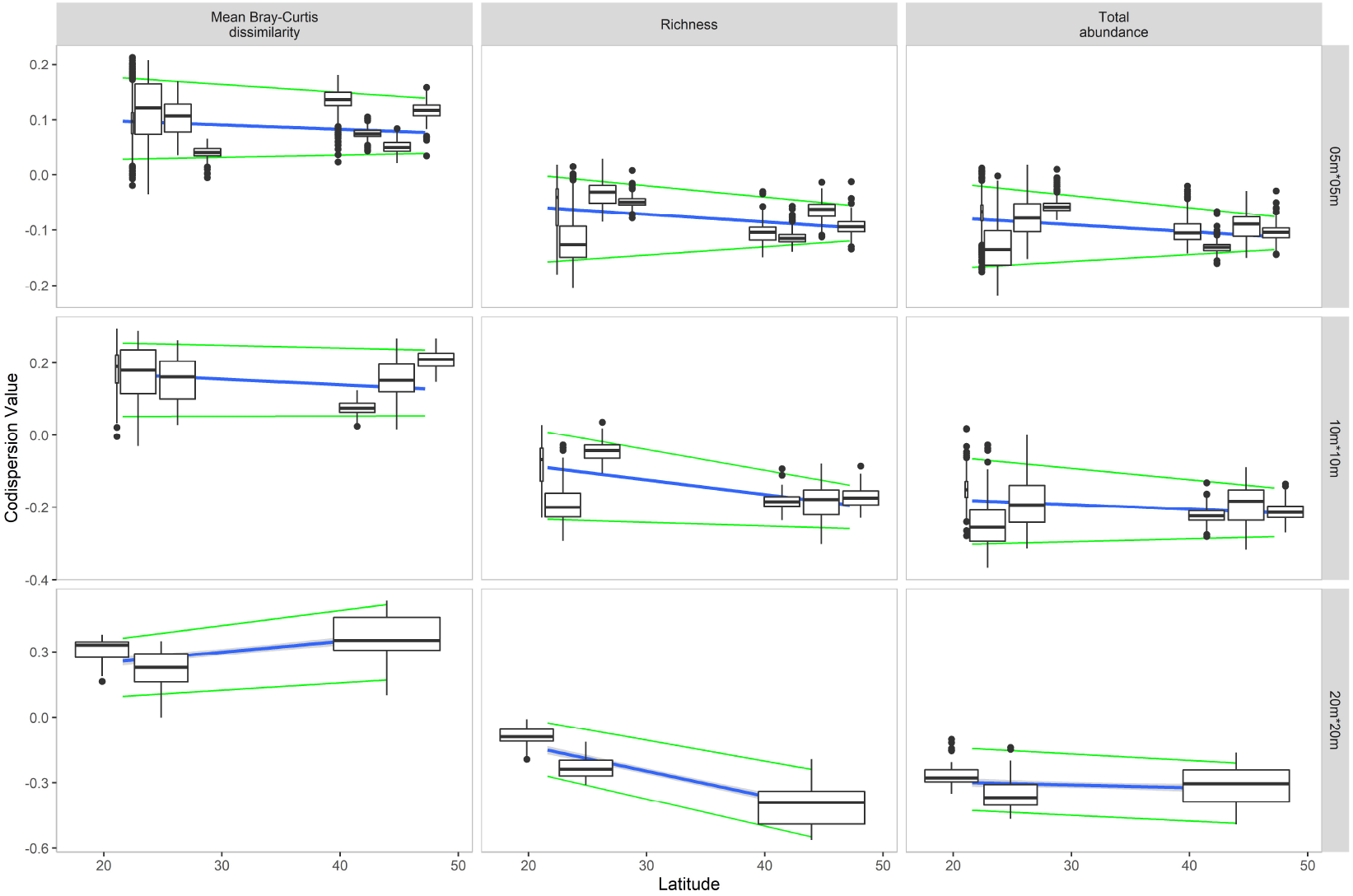
Relationship between latitude and codispersion between candidate foundation species (canopy trees and understory trees and shrubs) and three measures of associated woody-plant diversity at different spatial grains. Box plots illustrate median, upper and lower quartiles, and individual points outside of the upper and lower deciles of average codispersion at each latitude where candidate foundation species occurred (Table 3). Lines are regressions on all the data (blue lines), or on the 5% or 95% quantiles of the data (green lines).

**Figure 9:**
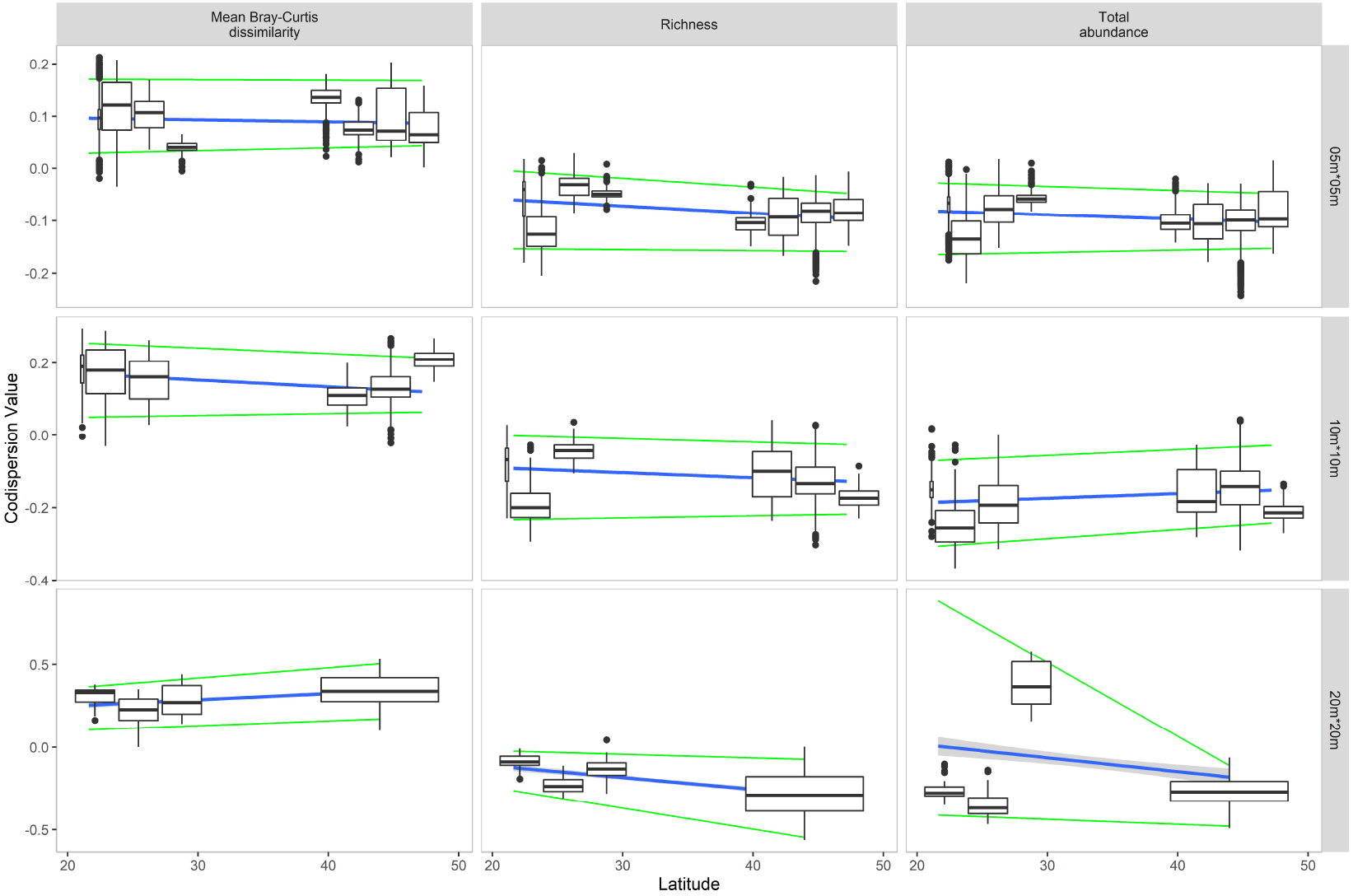
Relationship between latitude and codispersion between candidate foundation canopy tree species and three measures of associated woody-plant diversity at different spatial grains. Box plots illustrate median, upper and lower quartiles, and individual points outside of the upper and lower deciles of average codispersion at each latitude where candidate foundation species occurred (Table 3). Lines are regressions on all the data (blue lines), or on the 5% or 95% quantiles of the data (green lines).

## Discussion

We applied two new statistical criteria (Ellison et al., 2019) to screen 12 of the 17 CForBio Forest Dynamic plots in China for candidate foundation species. These 12 plots ranged from 47 to 21 *^◦^*N latitude, represented boreal, conifer-dominated, broad-leaved deciduous, subtropical, and tropical forests (Table 1), and included two forest types referred to by particular species (“Korean pine” mixed forests at Liangshi and Changbai Mountain, and the “*Taxus cuspidata*” mixed coniferous forest at Muling). Such eponyms do suggest traditional or cultural-based knowledge of foundation (or other “important”) species (Ellison et al., 2005; Ellison, 2019). Whilst both Korean pine (*Pinus koraiensis*) and *Taxus cuspidata* were identified as candidate foundation species (Table 3), they were only candidates in the Muling *Taxus cuspidata*-dominated forest plot, not in either of the “Korean pine” mixed forests. We also found a strong latitudinal gradient, unrelated to the expected (and observed) underlying latitudinal gradient in woody plant species richness, in the number of candidate foundation species, which were more frequent in temperate than in tropical forest plots (Fig. 7). Where they occurred, candidate foundation species had comparable effects at all latitudes (Figs. 8, 9), suggesting that foundation species effects more likely reflect specific combinations of traits and interspecific effects rather than being manifestations of “neutral” (sensu Hubbell, 2001) processes (Ellison et al., 2019).

### Candidate foundation species are more common in temperate latitudes

Foundation species in forests control species diversity locally within forest stands and at landscape and larger scales by creating habitat for associated flora (e.g., epiphylls, epiphytes, vines, lianas) and modifying soil structure and composition (e.g., Ellison et al., 2005; Brantley et al., 2013; Baiser et al., 2013; Vallejos et al., 2018; Degrassi et al., 2019; Ellison, 2019). Forest foundation species frequently are common, abundant, large trees (e.g., Schweitzer et al., 2004; Ellison et al., 2005; Whitham et al., 2006; Tomback et al., 2016; Ellison et al., 2019), but understory shrubs and treelets also can have foundational characteristics (Kane et al., 2011; Ellison and Degrassi, 2017; Ellison et al., 2019). Ellison et al. (2005) hypothesized that foundation species would be more likely in temperate forests because of their relatively low species richness and more frequent dominance by one or a small number of taxa. In contrast, tropical forests should lack foundation species as they are speciose and are dominated less frequently by a small number of taxa. Our data supported this hypothesis: candidate foundation species in the CForBio plots were more common at higher latitudes than in the tropics (Fig. 7; Ellison et al., 2019). This pattern also may reflect the greater importance of deterministic “niche” processes in temperate forests versus the stronger role of “neutral” dynamics in tropical ones (Gravel et al., 2006; Qiao et al., 2015).

We hypothesize that tropical forests dominated by a one or a few closely-related species, such as coastal mangrove forests dominated by *Rhizophora* spp. (Tomlinson, 1995) and monodominant tropical lowland forests dominated by species of Dipterocarpaceae in southeast Asia or species of Leguminosae (subfamily Caesalpinioideae) in Africa and the Neotropics (Torti et al., 2001; Hall et al., 2019) may be structured by foundation species (Ellison et al., 2005). Indeed, *Gilbertiodendron dewevrei* in the Ituri ForestGeo plot in the Democratic Republic of Congo (Makana et al., 2004*a*,*b*) has functional characteristics similar to *Tsuga canadensis* in northeastern US forests. *Gilbertiodendron* casts deep shade; produces leaf litter that decomposes very slowly, creating a dense and deep litter layer; creates soils with *≈*30% of the available nitrogen (ammonium + nitrate) relative to nearby mixed forests; and has a depauperate (albeit not unique) fauna of leaf-litter ants and mites (Torti et al., 2001). Analysis of species distribution and diversity associated with potential foundation species in Southeast Asian forests dominated by Dipterocarpaceae, such as the ForestGeo 50-ha Pasoh plot in Malaysia (Kochummen et al., 1991; Ashton et al., 2003) versus others lacking abundant dipterocarps, such as the 30-ha ForestGeo Mo Singto plot in Thailand (Brockelman et al., 2011) or the 2-ha plot in Aluoi, Vietnam (Nguyen et al., 2016) would provide useful comparisons with the analyses of the CForBio plots—especially the 20-ha Xishuangbanna plot—presented here.

Conversely, the mid-latitude peak in functional-trait diversity of trees (Lamanna et al., 2014) led Ellison et al. (2019) to hypothesize that foundation tree species should be less common in boreal forests at high latitudes or at high elevations in lower latitudes than in more temperate ones. Our data showing no candidate foundation species at the high-elevation but low-latitude Yulong Snow Mountain plot support this hypothesis (Table 3). In other high-elevation and high-latitude boreal ecosystems, foundation species tend to be low-growing perennial, cushion- or tussock-forming plants (e.g., Ellison and Degrassi, 2017; Elumeeva et al., 2017).

### Foundation species effects are scale-dependent at landscape, not local scales

Ellison (2019) argued that foundation species increase “patchiness” (beta diversity) at landscape scales, and that this effect of foundation species is of paramount importance when considering whether and how to conserve or otherwise manage them (see also Ellison et al., 2019). Across the 12 CForBio plots, we observed an increase in the strength of foundation species effects on beta diversity, expressed as a significant increase in codispersion between the candidate foundation species and diversity of associated species, at increasingly larger spatial grain (Fig. 6). At the 20-m grain, the magnitude of the codispersion coefficient approached that of many of the candidate foundation species in ForestGeo plots in the Americas (0.25–0.35; Fig. 6), but still less than the very strong effects of *T. canadensis* in northeastern US forests (Ellison et al., 2019).

Conversely, although foundation species can provide habitat for associated species, thus increasing their local diversity, the opposite pattern and magnitude of effects has been found when analyzing only associated woody plant species in forest dynamic plots (Buckley et al., 2016*a*; Ellison et al., 2019) because foundation species occupy most of the available space. In the CForBio plots, codispersion similarly was negative between candidate foundation species and alpha diversity of associated woody plants (Figs. 2–6), but this relationship did not vary significantly with spatial grain (Fig. 6). Additional data on faunal groups (e.g., Sackett et al., 2011; Record et al., 2018) or non-woody plants (e.g., Ellison et al., 2016) could provide a test of whether these candidate foundation species have a positive effect on other associated species that are not competing for space with canopy or subcanopy trees, but such data are collected rarely in forest dynamic plots (but see Schowalter, 1994; Ruchty et al., 2001; Ellison, 2018).

### *Acer* as a candidate foundation genus

In this study, four species of *Acer* were candidate foundation species among the three cold-temperate plots in China (Liangshi, Muling, and Changbai: Table 3). Among these, *A. ukurunduense* and *A. barbinerve* were the only two of all our candidate foundation species that met the most stringent criteria for consideration. In a comparable study across a latitudinal gradient in the Americas, *A. circinatum* was identified as a candidate foundation species in the the Wind River ForestGeo plot in Washington State, USA (Ellison et al., 2019). We hypothesize that in many forests throughout the Northen Hemishphere, that *Acer* not only can be a dominant genus in terms of abundance or total basal area, but that it may function as a foundation genus, akin to *Quercus* in the Tyson ForestGEO plot in central North America (Ellison et al., 2019).

*Acer* species often are common and abundant in temperate deciduous broad-leaved, coniferous, and mixed forests throughout the Holarctic (Tiffney, 1985; Pennington et al., 2004), and in subtropical montane forests in China (Xu, 1996). *Acer* includes ¿150 species (WFO (World Flora Online), 2020), at least 99 of which (including 61 endemics) occur in China (Xu et al., 2008) and more than a dozen are found in North America (Alden, 1995). *Acer* species generally are shade tolerant, (i.e., they can regenerate and grow under closed canopies) and have relatively high seedling and sapling survival rates (Tanaka et al., 2008). Some more shade-intolerant (“photophilous”) early-successional *Acer* species create conditions that facilitate restoration of both later successional forests and their associated animal assemblages (Zhang et al., 2010).

There are several forests named after *Acer* species in China, including the *Acer mono– Tilia amurensis–T. mandshurica* temperate broad-leaved deciduous forest, the *Schima superba– Acer caudatum–Toxicodendron succedaneum* eastern subtropical forest, and the *Cyclobalanopsis multinervis–Castanopsis eyrel* var. *caudata–Liquidambar acalycina–Acer sinense* forest in southwest China (Wu, 1995). *Acer* also are considered primary “companion” species in Chinese *Quercus* and mixed broad-leaved-Korean pine forests where multiple *Acer* species co-occur. For example, six–seven additional *Acer* species were recorded with the three candidate foundation *Acer* species in the two broad-leaved-Korean pine mixed forests plots (LS, CB). The nine *Acer* species in the CB plot account for ¿46% of the total stems (Zhang et al., 2010).

In North American forests, *Acer* species also define several forest types, including “Sugar Maple” (i.e., *A. saccharum*), “Sugar Maple –Beech–Yellow Birch”, “Sugar Maple–Basswood”, “Red Maple” (i.e., *A. rubrum*), and “Silver Maple–American Elm” (i.e., *A. saccharinum*) (Eyre, 1980). In forests of the Pacific Northwest of North America, the subcanopy treelet *A. circinatum* not only grows rapidly, has high biomass, and forms broad canopies that suppress other species (Lutz and Halpern, 2006; Halpern and Lutz, 2013), which causes it to have negative codispersion with other woody taxa (Ellison et al., 2019), but it also supports a high diversity of epiphytes (Ruchty et al., 2001). Another North American species, *A. saccharinum*, dominates floodplain forests on well-drained alluvial soils in the eastern U.S. (Gabriel, 1990). Although Vankat (1990) subsumed “Silver Maple–American Elm” forests within a “Mixed Hardwood Wetland Forest” type and considered *A. saccharinum* to be only a minor component of these forests, this species historically was a significant constituent of at least some primary forests in the upper Midwestern U.S. and Canada (Cho and Boerner, 1995; Simard and Bouchard, 1996; Guyon and Battaglia, 2018); supports unique assemblages of birds (Yetter et al., 1999; Knutson et al., 2005; Kirsch and Wellik, 2017); and, among woody species, contributes substantially to carbon fixation in tidal wetlands (Milligan et al., 2019). *Acer saccharinum* may be similar to other North American (candidate) foundation species whose effects are most pronounced at different successional stages (Ellison et al., 2014, 2019). However, we know of no large plots in either “Silver Maple–American Elm” or “Mixed Hard-wood Wetland” forests from which we could derive data to test whether *A. saccharinum* meets our statistical criteria for candidate foundation species. Whilst it may be premature to establish large forest dynamics plots in floodplains in either the temperate zone or the tropics, or in tropical coastal habitats with low tree diversity, comparable data could be used to test more general ideas about the foundational importance of particular genera, such as *Acer* or *Rhizphora*, in forested wetlands worldwide.

## Acknowledgments

This study was supported by the Strategic Priority Research Program of the Chinese Academy of Sciences (XDB31000000), the National Natural Science Foundation of China (31740551, 31270562, 31660130, 31760131), the National Key Research and Development Program of China (2016YFC0502405), the Guangxi Key Research and Development Program (AB17129009), Natural Science Foundation of Heilongjiang (CN) (QC2018025), and the Harvard Forest. We acknowledge the Chinese Forest Biodiversity Monitoring Network (CForBio) for installing and supporting the forest dynamic plots in China, and all the field technicians and students who have helped census the plots. Audrey Barker Plotkin, Brian Hall, Helen Murphy, Dave Orwig, Neil Pederson, and Jonathan Thompson provided constructive comments and feedback on early versions of the manuscript.

## Author contributions

XQ and AME conceptualized and designed the study and wrote the manuscript; XQ and all the other authors collected the data at the individual CForBio plots. All authors contributed critically to the drafts and gave final approval for publication.

## Supplementary Information

Table 4. Codispersion statistics for all candidate foundation species listed in Table 3.

